# Differential Patterns of Cross-Protection against Antigenically Distinct Variants in Small Animal Models of SARS-CoV-2 Infection

**DOI:** 10.1101/2024.03.14.584985

**Authors:** Prabhuanand Selvaraj, Charles B. Stauft, Shufeng Liu, Kotou Sangare, Tony T. Wang

## Abstract

Continuous evolution of severe acute respiratory syndrome coronavirus 2 (SARS-CoV-2) will likely force more future updates of vaccine composition. Based on a series of studies carried out in human ACE2 transgenic mice (K18-hACE2) and Syrian hamsters, we show that immunity at the respiratory tract, acquired through either previous infection or vaccination with an in-house live attenuate virus, offers protection against antigenically distinct variants in the absence of variant spike-specific neutralizing antibodies. Interestingly, immunity acquired through infection of a modern variant (XBB.1.5) was insufficient in preventing brain infection by the ancestral virus (WA1/2020) in K18-hACE2 mice. Similarly, previous infection with WA1/2020 did not protect against brain infection by XBB.1.5. Our results highlight the importance of immune components other than neutralizing antibodies in maintaining protection against new variants in the respiratory tract, but also paint scenarios where a monovalent vaccine based on a contemporary variant may be less effective against the ancestral strain.

**Importance:** Many studies have assessed the cross neutralization of various SARS-CoV-2 variants induced by breakthrough infections or vaccine boosters. Few studies, however, have modeled a more severe type of breakthrough infection. Here, we show that immunity acquired through a previous infection by either a historical virus (WA1/2020) or a contemporary variant (XBB.1.5) failed to protect against brain infection of K18-hACE2 mice by an antigenically distinct virus, although it largely protected the respiratory tract. Our results provided a potential model to investigate the role of different immune components in curbing SARS-CoV-2 infection.

## Introduction

Initial vaccination with two mRNA vaccines against SARS-CoV-2 provided up to 95% efficacy against severe disease outcomes (1, 2). The emergence of new SARS-CoV-2 variants, however, has significantly reduced vaccine effectiveness against symptomatic COVID-19. Since 2021, various Omicron sub-lineages have rapidly replaced previously circulating variants worldwide (3). Despite laboratory evidence of attenuated pathogenicity, increased transmissibility of Omicron sub-lineages led to large numbers of hospitalizations and deaths (4–6). New studies consistently demonstrate reduction in circulating neutralizing antibody (nAb) titers against emerging Omicron subvariants among those who received either the ancestral or bivalent (Wuhan-1 and BA.4/5) mRNA vaccines (7–10). For this reason, regulatory agencies from both the United States and Europe have recommended an updated monovalent XBB.1.5 vaccine composition.

A concern associated with a simplified vaccination regimen is that whether it offers protectivity against an earlier variant. In this study, we set out to test the cross protectivity offered by natural infection and vaccination with a live-attenuated vaccine candidate (LAV) against ancestral and contemporary SARS-CoV-2 variants in mice and Syrian hamsters. Our results show sustained protection in respiratory tracts in K18-hACE2 mice, but not in the brain. Vaccination with WA1/2020 (WA1) and BA.5 specific LAV candidates resulted in variant specific as well as cross-reactive humoral immunity and was protective in Syrian hamsters against EG.5.1 challenge. These results have significant implications in pre-clinical evaluation of future vaccines.

## Results

### Cross protection of convalescent K18-hACE2 mice in respiratory tract but not in brain in the absence of circulating nAbs

The K18-hACE2 mice are highly susceptible to SARS-CoV-2 with a lethal phenotype to certain variants (11–18). To derive convalescent animals, we infected K18-hACE2 mice with 100 plaque-forming units (PFU) of WA1 or the XBB.1.5 variant (Fig. 1A). Among those infected with WA1, around 20% of infected mice survived the infection. In contrast, more than half of the mice survived XBB.1.5 infection. While survivors experienced transient weight loss, they completely recovered their body weight and resumed healthy activity. At 60 days post infection (DPI), serum nAb titers were determined by 50% focus reduction neutralization test (FRNT_50_). All animals seroconverted and displayed significant nAb titers (measured by FRNT_50_) against the homotypic variant i.e., WA1 convalescent (anti [α]-WA1) [geometric mean titers (GMT) 787.2, interquartile range (IQR) 3631, p=0.0003] and XBB.1.5 convalescent (α-XBB.1.5) [GMT 174.7, IQR 1534, p=0.012] (Fig. 1B). By contrast, nAb titers of the WA1-convalescent K18-hACE2 mice were at or below limit of detection against XBB.1.5 [GMT 10, IQR 0.00]. Likewise, XBB.1.5-convalescent nAb titers against WA1 [GMT 11.33, IQR 47.36] were at or below the limit of detection. These results confirmed that mice infected with the ancestral or a contemporary variant of SARS-CoV-2 lacked nAbs against the antigenically distinct variant.

**Figure 1.**
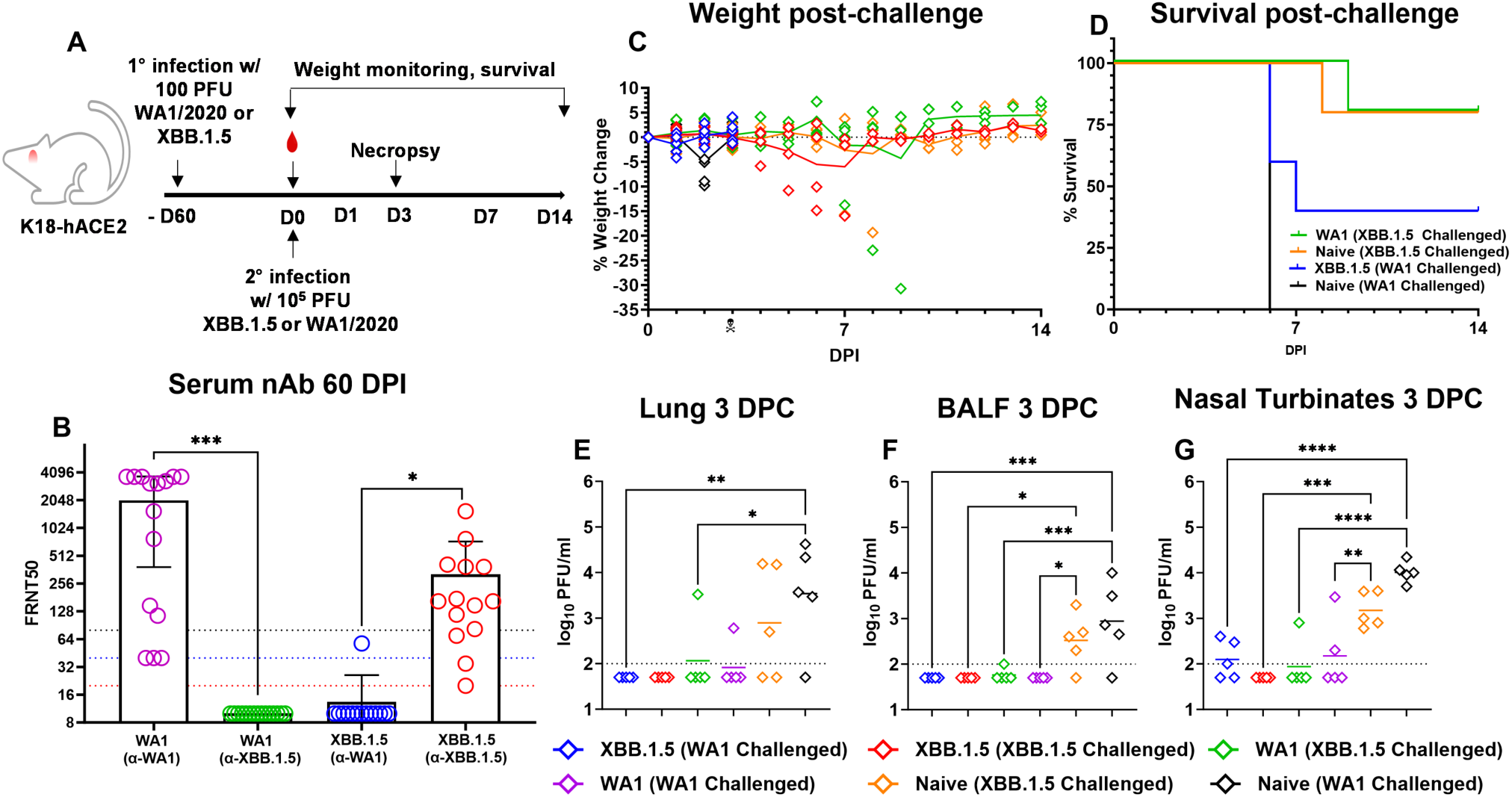
Convalescent K18-hACE2 mice are protected against rechallenge with a homologous or heterologous virus at respiratory tract. (**A**) Overall design of the study. (**B**) Serum collected at day 60 after the first low-dose infection (before rechallenging) tested for serum nAb against WA1 & XBB.1.5 in WA1 convalescent group (n=15 mice) or XBB.1.5 convalescent group (n=14 mice) by FRNT_50_ assay (paired t test; error bars indicate standard deviation; * p=0.012, *** p=0.0003). (**C**) Weight loss profile and survival (**D**) of immunized mice (WA1/2020 100 PFU or XBB.1.5 100 PFU) after a challenge with 10^5^ PFU of WA1 or XBB.1.5. (**E-G**) Lungs (*p=0.02, **p=0.004), BALF (*p=0.02, ***p=0.0005) and nasal turbinates (**p=0.008, ***p=0.0001, ****p<0.0001) were harvested at 3 DPC and measured for infectious viral titer by plaque forming assay analyzed using one-way analysis of variance (ANOVA) (n=5 animals each group).

Two months after the initial infection, we challenged convalescent and age-matched naive mice with 10^5^ PFU WA1 or XBB.1.5 to examine cross-protection in the absence of detectable nAbs (Fig. 1C-G). Four groups of mice (n= 10/group) were included: naïve mice challenged with WA1; naïve mice challenged with XBB.1.5; WA1 convalescent mice challenged with XBB.1.5; and XBB.1.5 convalescent mice challenged with WA1. After challenge, mice were monitored for weight loss and survival. Five mice from each group were euthanized on Day 3 after challenge for viral load analyses (for viral load analyses, two additional age-matched groups were included: convalescent mice that recovered from a previous WA1 infection were re-challenged with WA1; convalescent mice that recovered from a previous XBB.1.5 infection were re-challenged with XBB.1.5, Fig. 1E-G). At a dose of 10^5^ PFU, all naïve mice challenged with WA1 succumbed to infection by day 6. Around 50% of naïve mice challenged with XBB.1.5 had to be euthanized after meeting humane end points by day 7. Importantly, some convalescent mice that were challenged with a heterologous virus started losing weight on 6 DPC with 20% of XBB.1.5 (1 out of 5) and 60% of WA1 convalescent mice (3 out of 5) succumbing to infection by 9 DPC (Fig. 1C&D). Necropsy of the brain tissues collected post-mortem revealed evident viral infection in these animals (Fig. 2), confirming brain infection.

**Figure 2.**
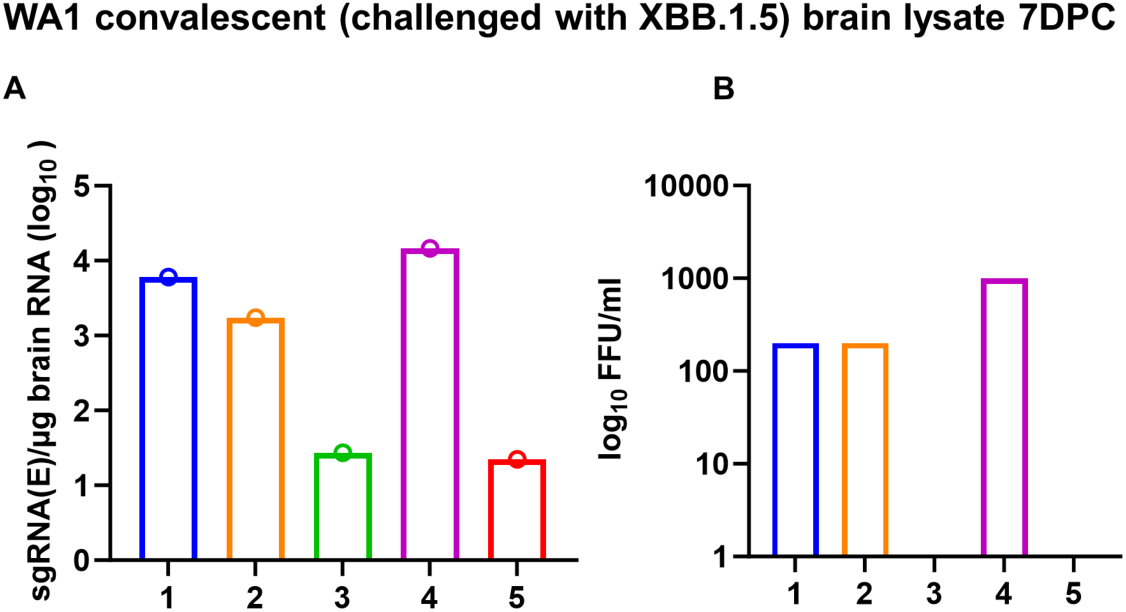
Presence of SARS-CoV-2 virus in K18-hACE2 mice that succumbed to infections. (**A**) Viral sgRNA levels in brains of WA1 convalescent (XBB.1.5 challenged) mice at 7 DPC were quantified by qRT-PCR. (**B**) Infectious virus from brain lysates were determined by focus-forming assays. Numbers 1-5 in panel A and B denote individual animals in the group.

In contrast to the observed lethality, at 3 DPC, convalescent mice largely showed undetectable viral load in the lungs [XBB.1.5 (WA1 Challenged) vs. Naive (WA1 Challenged), p=0.019; WA1 (XBB.1.5 Challenged) vs. Naive (WA1 Challenged), p=0.04] (Fig 1E). Similar undetectable viral load levels were found in bronchoalveolar lavage fluid (BALF) [XBB.1.5 (WA1 Challenged) vs. Naive (WA1 Challenged), p=0.001; XBB.1.5 (XBB.1.5 Challenged) vs. Naive (XBB.1.5 Challenged), p=0.01; WA1 (XBB.1.5 Challenged) vs. Naive (WA1 Challenged), p=0.001] (Fig 1F).Sporadic infectious viral load were detected in nasal turbinates of convalescent mice but levels remained significantly lower than naïve mice [XBB.1.5 (WA1 Challenged) vs. Naive (WA1 Challenged), p<0.0001; XBB.1.5 (XBB.1.5 Challenged) vs. Naive (XBB.1.5 Challenged), p=0.0001; WA1 (XBB.1.5 Challenged) vs. Naive (WA1 Challenged), p=0.0001; WA1 (WA1 Challenged) vs. Naive (XBB.1.5 Challenged), p=0.007] (Fig. 1G). Notably, convalescent mice that received a heterologous viral challenge demonstrated similar viral loads in the lung compared to convalescent mice that received a homologous viral challenge, despite the absence of nAbs for the challenging strain. These results indicate that natural immunity obtained from previous infection remains cross-protective in the lower respiratory tract against challenge with a distant SARS-CoV-2 variant.

We subsequently performed hematoxylin and eosin (H&E) staining on lung tissues collected at 3 DPC. Histopathology was found to correlate with viral loads (Fig. 3A-L). While naïve mice developed observable pathologies, including alveolar wall thickening and alveolar airway infiltrates in the lungs after virus challenge, convalescent mice showed fewer histological changes after challenge with either a homologous or a heterologous virus (Fig. 3M-N). We also assessed immune activation by immunohistochemistry and found CD4^+^ and CD8^+^T cells in the lungs of all groups irrespective of convalescent or homologously- or heterologously challenged mice at 3 DPC (Fig. 4).

**Figure 3.**
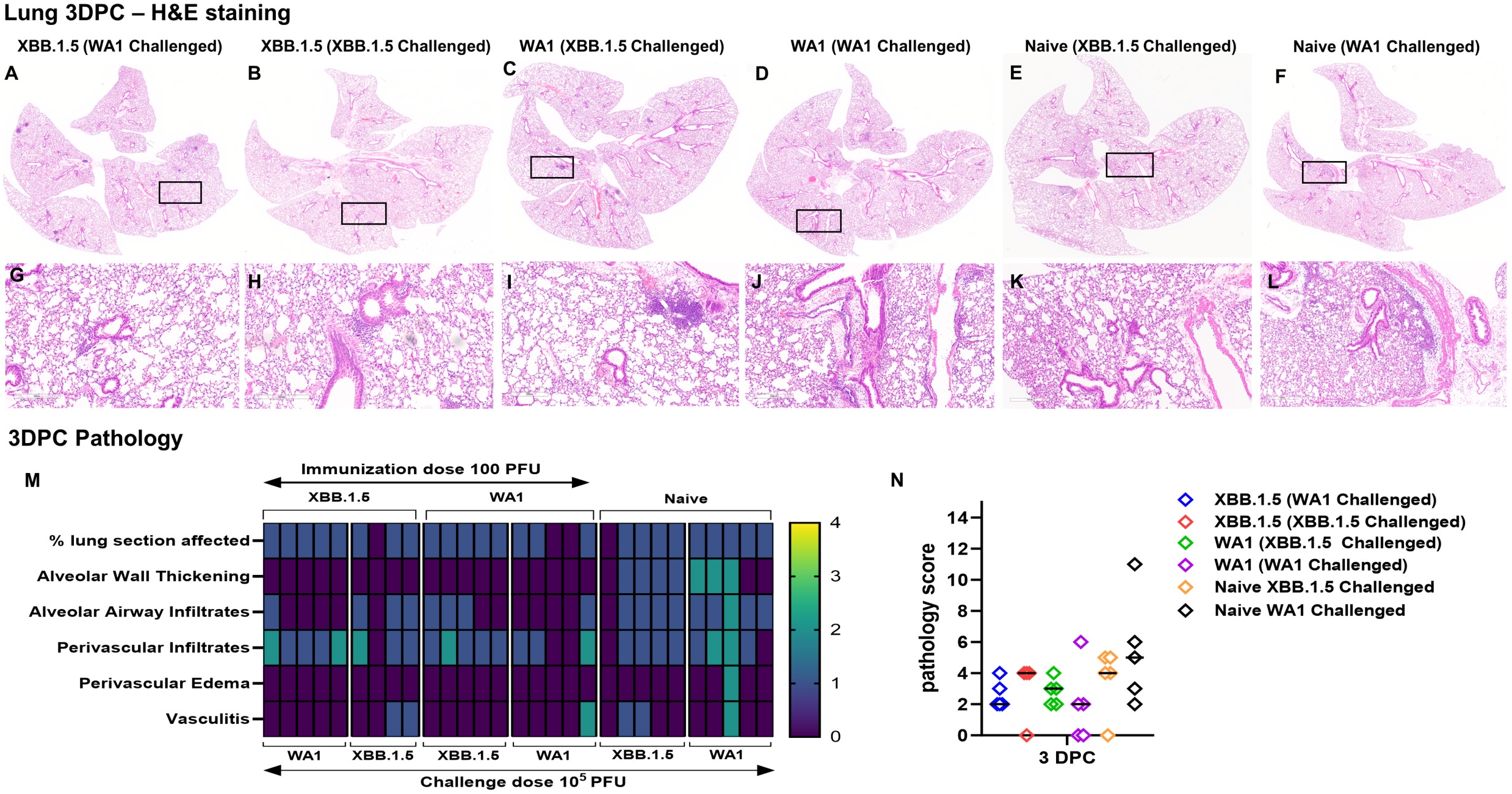
Examination of histopathology corroborates cross protection of convalescent K18-hACE2 mice against homologous or heterologous virus challenges. (**A-L**) Representative images of H&E-stained lung tissue sections (5 µm thickness) collected at 3 DPC. (**A, G**) XBB.1.5 convalescent (WA1 challenged), (**B, H**) XBB.1.5 convalescent (XBB.1.5 challenged), (**C, I**) WA1 convalescent (XBB.1.5 challenged), (**D, J**) WA1 convalescent (WA1 challenged), (**E, K**) naïve (XBB.1.5 challenged), (**F, L**) naïve (WA1 challenged). Black rectangular boxes in (**A**), (**B**), (**C**), (**D**), (**E**) indicate regions from which magnified images in (**G**), (**H**), (**I**), (**J**), (**K**) and (**L**) are taken. Scale bars in (**A-F**) 4mm and 0.6x magnification, (**G-L**) 300µm & 8.8x magnification. (**M**) Heatmap presentation of histopathology scores of lungs collected at 3 DPC based on each category (refer method section for scoring criteria). (**N**) Cumulative histopathology scores of infected lungs at 3 DPC (n=5 mice in all groups except (n=4) in XBB.1.5 convalescent (XBB.1.5 challenged) group. Bars in the scatter dot plot indicate median values.

**Figure 4.**
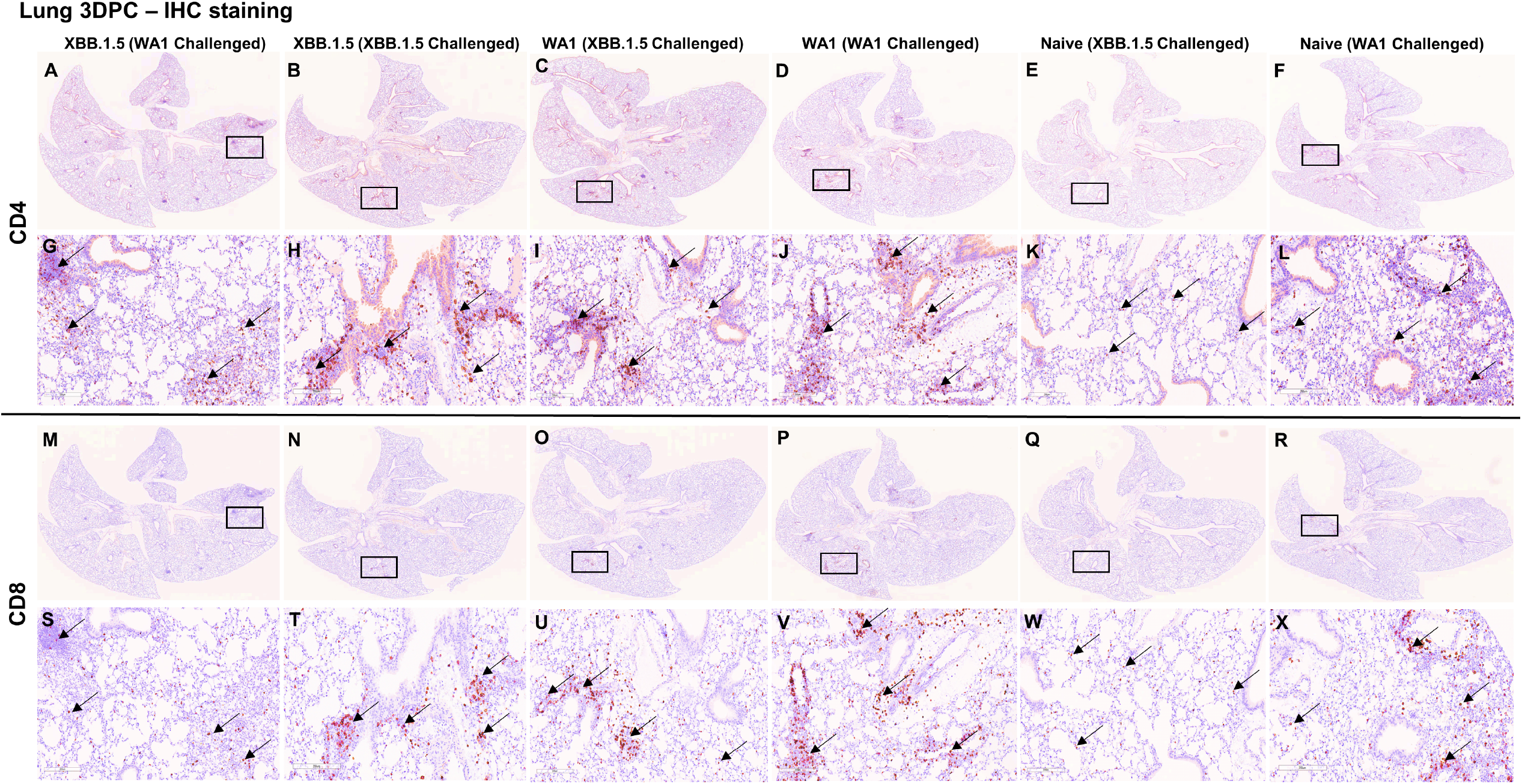
Immune activation in the lungs. Lung tissue sections from 3 days post challenged mice were subjected to immunohistochemistry staining for CD4 and CD8 T cell markers. Panel (**A**-**L**) represents CD4 T cell marker staining and (**M**-**X**) represents CD8 T cell marker staining. (**A, G, M, S**) XBB.1.5 convalescent (WA1 challenged), (**B, H, N, T**) XBB.1.5 convalescent (XBB.1.5 challenged), (**C, I, O, U**) WA1 convalescent (XBB.1.5 challenged), (**D, J, P, V**) WA1 convalescent (WA1 challenged), (**E, K, Q, W**) naïve mice XBB.1.5 challenged, (**F, L, R, X**) naïve mice WA1 challenged. Black rectangular boxes indicate magnified regions. (**G**), (**H**), (**I**), (**J**), (**K**), (**L**) are closeup images of (**A**), (**B**), (**C**), (**D**), (**E**), (**F**), and (**S**), (**T**), (**U**), (**V**), (**W**), (**X**) are closeup images of (**M**), (**N**), (**O**), (**P**), (**Q**), (**R**) respectively. Black arrows that point to brown staining in panels (**G**-**L**) indicate CD4 positive staining and in panels (**S**-**X**) indicate CD8 positive staining. Scale bars in (**A-F** & **M-R**) 4mm and 0.6x magnification, (**G-L** & **S-X**) 200µm & 13.2x magnification.

### Cross protectivity conferred by candidate live attenuated vaccine viruses

Our laboratory has previously designed multiple LAV candidates in which three attenuating modifications were introduced into the WA1 genome: the removal of the furin cleavage site, the deletion of ORFs 6-8, and introduction of a pair of mutations to the Nsp1 gene (21). To determine if a vaccine designed against an ancestral virus may protect against contemporary variants, we tested our LAVs in Syrian hamsters, which are highly susceptible to SARS-CoV-2 and have been widely used in COVID-19 research (23,(19). Besides the prototype WA1-based LAV, we also generated additional attenuated viruses with the WA1 spike replaced with BA.1, BA.2 or BA.5 spike protein, namely BA.1-LAV, BA.2-LAV, and BA.5-LAV (23). We vaccinated male adult Syrian hamsters with a single, low dose (100 PFU) of LAVs bearing WA1 spike, BA.5 spike or mixed LAV (WA1 + BA.1 + BA.2 + BA.5) (Fig.5A). At 22- and 225-days post vaccination, serum nAb titers against XBB.1.5 (Fig. 5B), EG.5.1 (Fig. 5C), and WA1 (Fig. 5D) were determined from each hamster by FRNT_50_ assay. As expected, of the WA1-LAV vaccinated hamsters (n=8), only 12.5% (n=1) had nAb titers against XBB.1.5 (GMT 13.5) and 50% (n=4) against EG.5.1 (GMT 21.9) at 225 DPI. Of the BA.5-LAV vaccinated hamsters, 62.5% had detectable anti-XBB.1.5 (GMT 21.8) nAbs and 100% (n=8) had nAbs against EG.5.1 (GMT 37.9) at 225 DPI. Interestingly, although the BA.5-LAV vaccinated group did not initially have detectable nAbs against WA1, all animals seroconverted by 225 DPI (∼4-fold increase in anti-WA1 nAb titers). Of the LAV-Mix vaccinated hamsters (n=4), all developed nAB antibodies against XBB.1.5 (Fig 5B) and EG.5.1 (Fig 5C). Hamsters receiving the mixture of attenuated viruses had the highest geometric mean nAb titers against XBB.1.5 (GMT 63.2) and EG.5.1 (GMT 70.1) at 22 DPI (Fig 5B&C).

**Figure 5.**
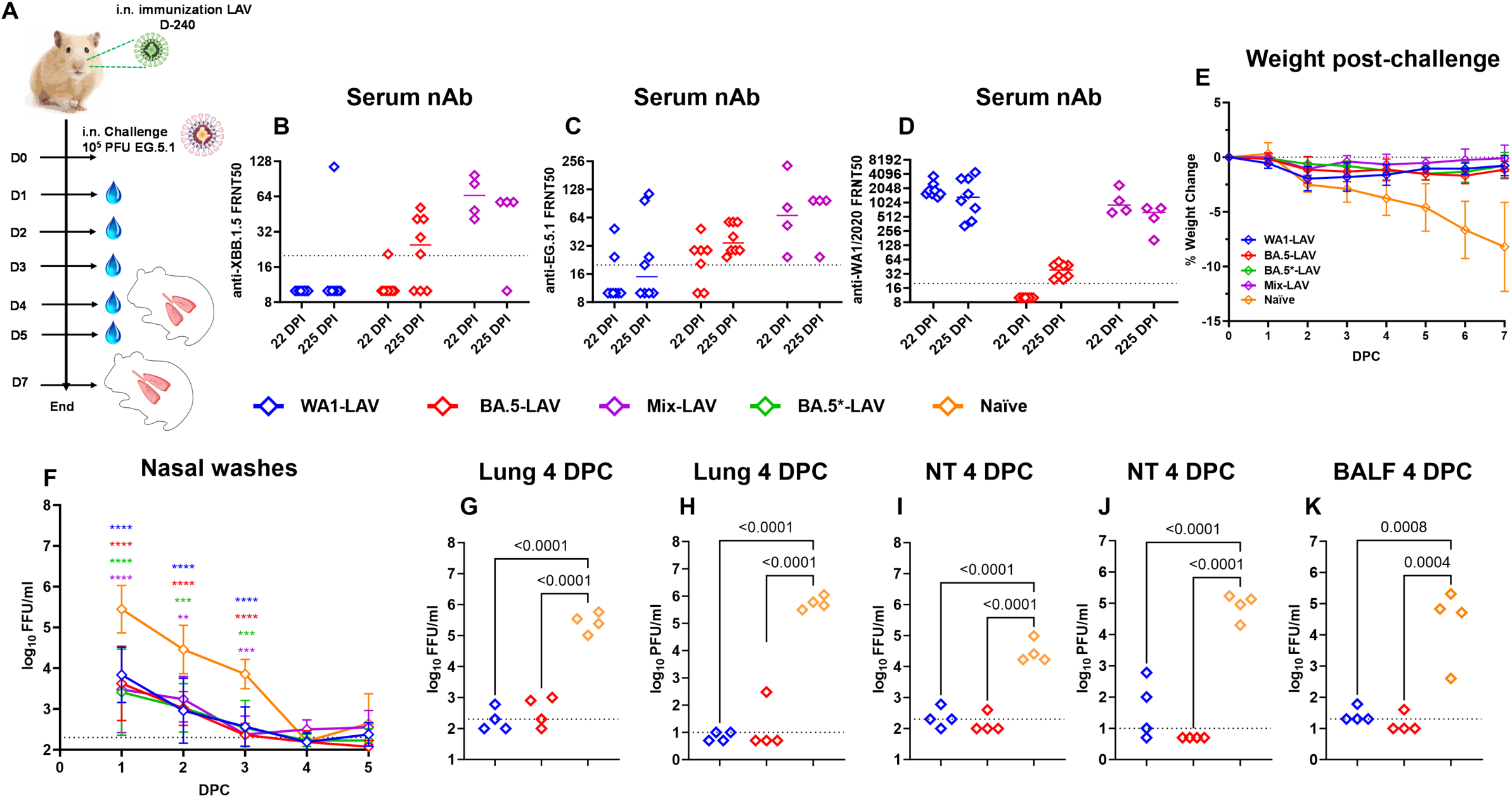
Vaccinations of LAV candidates protect Syrian hamster against EG.5.1 challenge. **(A)** Overall design of the study. Three months old male Syrian hamsters were intranasally inoculated with LAVs (100 PFU Nsp1-WA1; blue; n=8 or Nsp1-BA.5; red; n=8; or 100 PFU concoction mixture [25 PFU Nsp1-WA1 + 25 PFU Nsp1-BA.1 + 25 PFU Nsp1-BA.2 +25 PFU Nsp1-BA.5]; violet; n=4 or Nsp1-BA.5* star indicates 10^4^ PFU; green; n=4;) and naïve (brown; n=8) group hamster with PBS. Bars in scatter dot plot bar indicate median values. Animals were bled at 22- and 225-days post immunization to collect sera and tested for circulating neutralizing antibodies against XBB.1.5 (**B**), EG.5.1 (**C**) and WA1 (**D**) by FRNT_50_ assay. (**E**) Weight change recorded for 7 DPC with 10^5^ PFU EG.5.1. (**F**) Nasal wash viral titer detected by focus forming assay (p**=0.0018, p***=0.0002, p****<0.0001; 2-way ANOVA Dunnett’s multiple comparison test). Infectious viral titers from lung homogenates, nasal turbinates and BALF at 4 DPC were determined by focus forming assay (**G**, **I**, **K**) or plaque forming assay (**H**, **J**); n=4 animals in each group.

On day 241 after the initial vaccination, WA1-LAV, BA.5-LAV, BA.5* LAV (vaccinated with 10^4^ PFU) and Mix-LAV vaccinated hamsters were challenged with 10^5^ PFU of the EG.5.1 variant. Vaccinated hamsters had minimal (<2%) body weight loss over 7 days-post-challenge (Fig. 5E). By contrast, naïve (unvaccinated) hamsters had up to 12% body weight loss at day 7 following challenge. The overall amounts of infectious virus measured from nasal washes were comparable amongst the three vaccinated groups, which were 10-100-fold lower than that from the naïve group at 1 DPC (WA1-LAV, BA5-LAV, BA5*-LAV and Mix-LAV, p<0.0001), 2 DPC (WA1-LAV and BA5-LAV, p<0.0001; BA5*-LAV, p=0.0002 and Mix-LAV, p=0.0018) and 3 DPC (WA1-LAV and BA5-LAV, p<0.0001; BA5*-LAV and Mix-LAV, p=0.0001) (Fig. 5F). Infectious virus titers in the lungs, nasal turbinates, and BALF at 4 DPC were also quantified using focus and plaque forming assay (Fig. 5G-K). Compared to unvaccinated naïve mice, all vaccinated animals had 100-1000-fold reduction in viral loads in the lungs (Fig 5G&H), nasal turbinates (WA1-LAV, BA5-LAV, p<0.0001) (Fig 5I&J), and in BALF (WA1-LAV p=0.0008, BA5-LAV p=0.0004) (Fig 5K). Generally, infectious viral loads from vaccinated hamsters were near, at, or below the limit of qualification of the assay, indicating that a robust protection against a contemporary variant was achieved in the respiratory tract after LAV vaccination.

To corroborate the above observation, we also performed H&E staining of the hamster lungs (Fig.6). For this part of the study, BA.5*-LAV and Mix-LAV vaccinated hamsters (n=4) were euthanized at 7 DPC while animals in the WA1-LAV, BA.5-LAV and naive groups were euthanized at both 4 DPC and 7 DPC (n=4). EG.5.1 infection induced mild to moderate levels of lung consolidation in naïve (unvaccinated) animals (Fig. 6A). Other observable pathologies included alveolar wall thickening, alveolar airway infiltration, perivascular infiltrates, alveolar osteoid foci, bronchiole mucosal hyperplasia, bronchiole airway infiltrates, protein fluid and vasculitis, bronchiolar necrosis, alveolar metaplasia, and atypical pneumocyte hyperplasia. By contrast, the integrity of alveolar space was preserved among all LAV vaccinated hamster groups after EG.5.1 challenge (Fig.6A-S). Compared to the naïve group, WA1-LAV, BA.5-LAV and Mix-LAV vaccination reduced total pathology scores with higher dose of BA.5-LAV (BA.5*-LAV group) vaccination being slightly more effective at 7DPC (Fig. 6T). Together, immunization of male Syrian hamsters with our LAV candidates protected against an antigenically distal variant in the lung.

**Figure 6.**
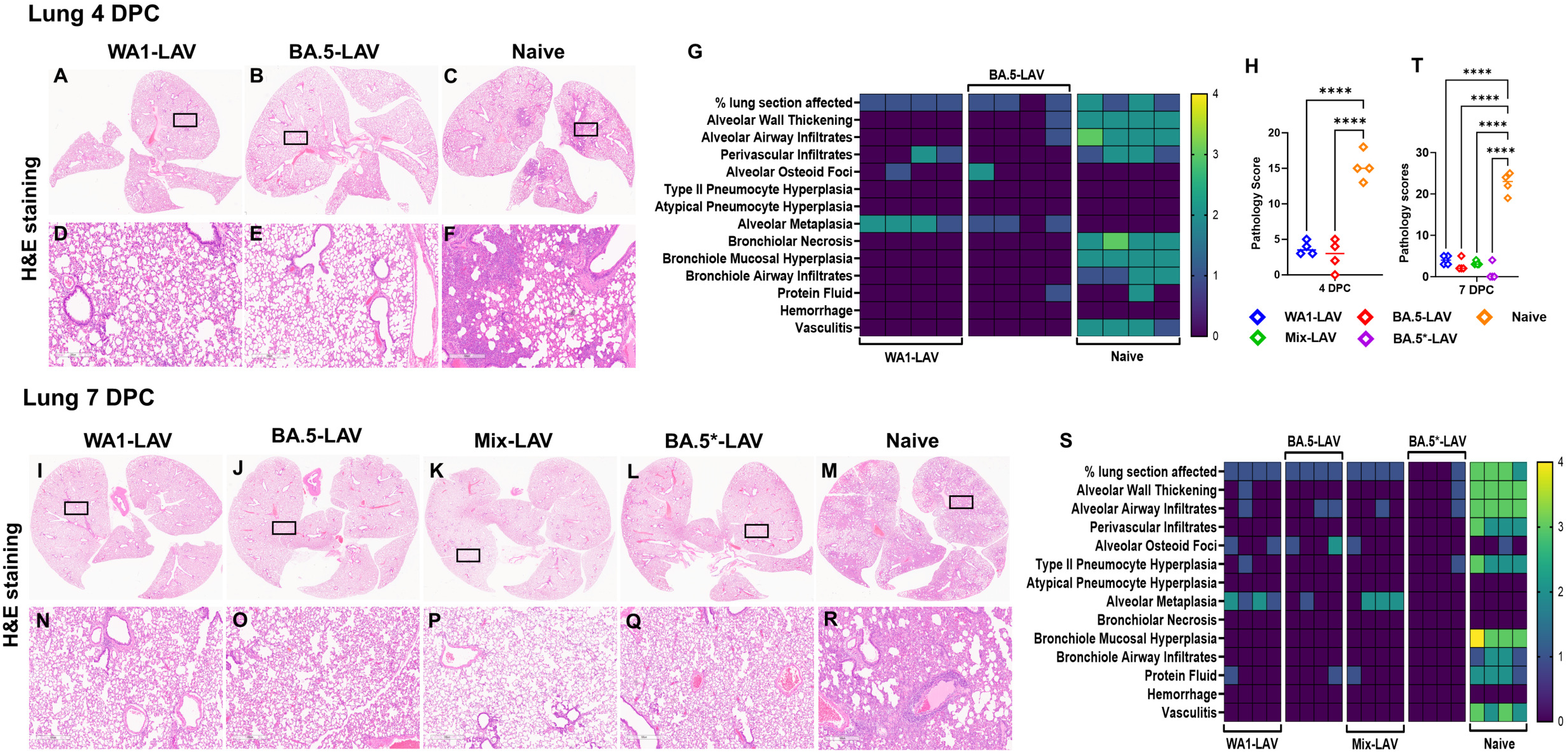
Significant reduction of lung pathology in LAV-vaccinated hamsters upon EG.5.1 challenge. Representative images of H&E-stained lungs from LAV immunized hamsters challenged with 10^5^ PFU EG.5.1 (**A**-**F**) from 4 DPC and (**I**-**R**) 7 DPC. Rectangular black box in (**A**-**C**) and (**I**-**M**) image indicates magnified region. (**D**), (**E**), and (**F**) are closeup image of (**A**), (**B**) and (**C**) respectively. Similarly (**N**), (**O**), (**P**), (**Q**) and (**R**) are closeup image of (**I**), (**J**), (**K**), (**L**) and (**M**) respectively. Scale bar = 6mm (**A**-**C**, **I**-**M**) and 500µm (**D**-**F**, **N**-**R**). (**G**) 4 DPC and (**S**) 7 DPC heatmap of histopathology scores in the lungs plotted based on each category (refer methods section for criteria details). (**H**) and (**T**) are cumulative lung pathology scores of 4 DPC and 7 DPC. Rhombus symbols in the scatter dot plot: blue (Nsp1-WA1), red (Nsp1-BA.5), green (Nsp1-Mix), violet (Nsp1-BA.5*) and brown (naïve). Experiment used n=4 animals per group.

## Discussion

Although the Omicron lineage of SARS-CoV-2 variants might exhibit reduced pathogenicity, the number of hospitalization of patients and cases have not been reduced (20, 21). Additionally, SARS-CoV-2 continues to evolve to escape the population immunity elicited by either natural infection or vaccination. For these reasons, research and preventive measures have been focused on how to improve the available vaccines to combat new variants. Because of so many breakthrough infections in humans, it is postulated that immunity acquired from a previous infection is insufficient to control reinfection with an antigenically distinct variant. We tested such a possibility by infecting K18-hACE2 mice with WA1 or XBB.1.5 and rechallenging them with a higher dose of homologous or heterologous virus. Our results show that immunity developed against previous SARS-CoV-2 infection offers protection against reinfection irrespective of SARS-CoV-2 variants in the respiratory tract of K18-hACE2 mice, even in the absence of variant spike-specific neutralizing antibodies. This finding implies that non-neutralizing antibodies and/or cellular immunity effectively offer cross protection against the tested SARS-CoV-2 variant in our experimental model. Differing from what was observed in respiratory tract, a fraction of convalescent K18-hACE2 mice still developed viral infection in the brain upon heterologous virus challenge and ultimately succumbed to infection. The lethality of SARS-CoV-2 in K18-hACE2 mice is known to be caused by encephalitis (11–17). Such a contrast between respiratory tract and brain suggests that neutralizing antibodies may be necessary to block the virus from invading the central nervous system in mice.

To expand the above observations into vaccine research, we chose an in-house developed LAV candidate to induce immunity on par with a natural infection. In this study, we show sustained (∼8 month) protection against EG.5.1 challenge in respiratory tract in Syrian hamsters after vaccination with a LAV based on the ancestral SARS-CoV-2 isolate WA1. Given the significant antigenic difference between WA1 and EG.5.1, our results demonstrate that LAVs can be an attractive addition to the panoply of vaccination strategies in place to combat the ongoing COVID-19 pandemic. Near-universal population immunity to SARS-CoV-2 may somewhat mitigate the safety concerns of using LAV in immune-competent individuals as a booster against emerging variants. In support, we have shown previously (22, 23) that LAV can boost heterotypic nAbs in previously vaccinated hamsters. Similarly, another research group in Syrian hamster model with an LAV candidate (sCPD9) also shown vaccinated animals developing robust nAbs against heterologous SARS-CoV-2 challenge (24). This LAV strategy could mitigate concerns over specific antigen-based, vaccine strategies, which can lose protection over time and may lack protection against emerging variants. Notably, mixing LAVs (WA1 + BA.1 + BA.2 + BA.5) induced higher nAB titers against EG.5.1 than did a single LAV, implying that a mixture of antigens might induce a more diverse B cell antibody repertoire, which cross-neutralize a future new variant.

There are a few limitations to this study. Due to the constant evolving nature of SARS-CoV-2, this study was not carried with the most recent circulating SARS-CoV-2 variant (i.e., JN.1). The observed brain infection of K18-hACE2 mice during a heterologous virus challenge may not be extrapolated to humans because of the evident anatomic and immunological differences between the two species. Nonetheless, profiling other immune markers in K18-hACE2 mouse model would likely shed lights on how the host manages the burden of immune evasive variants without specific neutralizing antibodies.

## ACKNOWLEDGEMENTS

The following reagent was deposited by the Centers for Disease Control and Prevention and obtained through BEI Resources, NIAID, NIH: SARS-Related Coronavirus 2, Isolate hCoV-19/USA-WA1/2020, NR-52281. The following reagents were obtained through BEI Resources, NIAID, NIH: SARS-Related Coronavirus 2, Isolate hCoV-19/USA/MD-HP47946/2023 (EG.5.1) NR-59576 and hCoV-19/USA/MD-HP40900/2022 (XBB.1.5) NR-59105, contributed by Andrew S. Pekosz. We are grateful to Dr. P. Shi (UTMB) for the original 7 plasmids to perform SARS-CoV-2 reverse genetics. We thank Dr. C. Florence for assistance in obtaining critical reagents and Dr. A. Soare for critical reading of the manuscript. The work described in this manuscript was supported by U.S. FDA intramural grant funds. The funders had no role in study design, data collection and analysis, decision to publish, or preparation of the manuscript.

## Materials and Methods

### Cells and Viruses

Vero E6 cell line (Cat # CRL-1586) was purchased from American Type Cell Collection (ATCC) and cultured in eagle’s minimal essential medium (MEM) supplemented with 10% fetal bovine serum (Invitrogen) and 1% penicillin/streptomycin and L-glutamine. A549-hACE2 (NR-53821) cells were obtained from BEI Resources and maintained in DMEM supplemented with 5% penicillin and streptomycin, and 10% fetal bovine serum at 37 °C with 5% CO2. H1299-hACE2 is a human lung carcinoma cell line stably expressing human ACE2. The cell line was generated by lentiviral transduction of the NCI-1299 human lung carcinoma cell line (ATCC CRL-5803) with pLVX-hACE2 and selected with 1 μg/mL puromycin. A western blot was performed to confirm the expression of hACE2. H1299-hACE2 cells were maintained in DMEM supplemented with 5% penicillin and streptomycin, and 10% fetal bovine serum at 37 °C with 5% CO_2_.

The SARS-CoV-2 isolate WA1/2020 (NR-52281, lot 70033175), XBB.1.5 (hCoV-19/USA/MD-HP40900/2022, NR-59105), and EG.5.1 (hCoV-19/USA/MD-HP47946/2023, NR-59576) were obtained from BEI Resources, NIAID, NIH. WA1/2020 had been passed three times on Vero cells and 1 time on Vero E6 cells prior to acquisition. It was further passed once on Vero E6 cells in our lab. The virus has been sequenced and verified to contain no mutation to its original seed virus. XBB.1.5 and EG.5.1 were used immediately upon receiving without subsequent passages. Generation of live attenuated viruses has been described elsewhere (22, 23).

### Mouse Infection Experiments

Female adult K18-hACE2 mice were previously purchased from the Jackson laboratory and held at FDA vivarium. All experiments were performed within the biosafety level 3 (BSL-3) suite on the White Oak campus of the U.S. Food and Drug Administration. The study protocol details were approved by the White Oak Consolidated Animal Care and Use Committee and carried out in accordance with the PHS Policy on Humane Care & Use of Laboratory Animals.

For infection studies, mice (7-8 weeks old) were first anesthetized by 3-5% isoflurane. Intranasal inoculation was done by pipetting 100 PFU (WA1 or XBB.1.5) SARS-CoV-2 in 25 µl volume dropwise into the nostrils of the mouse. Mice were weighed and observed daily. After 6 weeks post infection, mice were bled to collect serum samples and stored at −80°C for future use to determine circulating neutralizing antibody titers against WA1 or XBB.1.5. At day 60 mice were rechallenged with 10^5^ pfu of WA1 or XBB.1.5, were weighed and monitored for any clinical signs of sickness. For tissues and bronchoalveolar lavage fluid (BALF) collection mice were euthanized by CO_2_ overdose on day 3 post challenge as necessary.

### Hamster Challenge Experiments

Male outbred Syrian hamsters (Mesocricetus auratus) were previously purchased from Envigo and held at FDA vivarium. All experiments were performed within the biosafety level 3 (BSL-3) suite on the White Oak campus of the U.S. Food and Drug Administration. The animals were implanted subcutaneously with IPTT-300 transponders (BMDS), randomized, and housed 2 per cage in sealed, individually ventilated rat cages (Allentown). Hamsters were fed irradiated 5P76 (Lab Diet) *ad lib*, housed on autoclaved aspen chip bedding with reverse osmosis-treated water provided in bottles, and all animals were acclimatized at the BSL3 facility for 4-6 days or more prior to the experiments. The study protocol details were approved by the White Oak Consolidated Animal Care and Use Committee and carried out in accordance with the PHS Policy on Humane Care & Use of Laboratory Animals.

For vaccination studies, three-month-old Syrian hamsters were anesthetized with 3-5% isoflurane following procedures as described previously (24, 25). Hamsters were intranasal inoculated by pipetting 100 PFU of attenuated viruses having WA1 spike or BA.5 spike or 10^4^ PFU of attenuated virus with BA.5 spike or inoculated with 25 PFU each of attenuated viruses having WA1 spike, BA.1 spike, BA.2 spike, BA.5 spike (100 PFU total in the final mixture) in 50 µl volume dropwise into the nostrils of the hamster under anesthesia. Nasal washes were collected by pipetting ∼160 µl sterile phosphate buffered saline into one nostril when hamsters were anesthetized by 3-5% isoflurane. Eight months later hamsters were challenged with 10^5^ PFU of EG.5.1 in 50 µl volume dropwise into nostrils, were weighed and monitored for clinical signs of sickness. For tissues and BALF collection, a subset of hamsters was humanely euthanized by intraperitoneal injection of pentobarbital at 200mg/kg and lungs were harvested for histopathology. Blood collection was performed under anesthesia (3-5% isoflurane) through gingival vein puncture.

### Virus titration

Nasal wash, BALF, and lung homogenate samples were 10-fold serially diluted in 96-well plates and dilutions added to 96-well black-well plates for fluorescent focus forming assays in H1299-hACE2 cells (26). After 1 hour incubation at 37°C, the Tragacanth gum overlay (final concentration 0.3%) was added. Cells were incubated at 37°C and 5% CO_2_ overnight (∼18 hours), then fixed with 4% paraformaldehyde, followed by primary staining of cells with rabbit anti-N Wuhan-1 antibody (Genscript U739BGB150-5) (1:2000 dilution) overnight. Cells were then washed three times with PBS 0.1% Tween-20, followed by secondary staining with anti-rabbit Alexa-488 conjugated antibody (1:2000 dilution), and 4′,6-diamidino-2-phenylindole (DAPI). The infectious titers were then counted using Gen5 software on a Cytation7 (Agilent) and calculated and plotted as focus forming units per milliliter (FFU/ml). Plaque forming assays in Vero E6 cells were performed as described previously (Liu et al., 2022).

### Real-time PCR assay of SARS-CoV-2 subgenomic RNA

Quantification of SARS-CoV-2 E gene subgenomic mRNA (sgmRNA) was conducted using Luna Universal Probe One-Step RT-qPCR Kit (New England Biolabs) on a Step One Plus Real-Time PCR system (Applied Biosystems). The primer and probe sequences were: SARS2EF: CGATCTCTTGTAGATCTGTTCT; PROBE: FAM-ACACTAGCCATCCTTACTGCGCTTCG-BHQ-1; SARS2ER: ATATTGCAGCAGTACGCACACA. To generate a standard curve, the cDNA of SARS-CoV-2 E gene sgmRNA was cloned into a pCR2.1-TOPO plasmid. The copy number of sgmRNA was calculated by comparing to a standard curve obtained with serial dilutions of the standard plasmid. The limit of quantification (LoQ) of the sgmRNA was determined to be 100 copies/reaction. Values below LoQ were mathematically extrapolated based on the standard curves for graphing purpose. When graphing the results in Prism 10, values below LoQ were arbitrarily set to half of the LoQ values.

### SARS-CoV-2 neutralization assay

Samples were serially diluted 2-fold in 5% FBS DMEM and mixed with 100-200 FFU of SARS-CoV-2 (WA1 or XBB.1.5 or EG.5.1) in a 96-well plate at 37°C for 1 hour. Sample: virus mixtures were then added to confluent H1299-hACE2 cells in 96-well plates. Cells were infected for 1 hour before the inoculum was removed and washed three times with DPBS. A second overlay containing 1.2% Tragacanth gum, 2X MEM, 5% FBS, and DMEM was added to the plate. Cells were incubated at 37°C for 1 day, then fixed with 4% paraformaldehyde, followed by staining of cells with primary rabbit anti-SARS-CoV-2 N Wuhan-1 antibody (Genscript U739BGB150-5) (1:2000 dilution) overnight followed by secondary anti-rabbit Alexa-488 conjugated antibody (1:2000 dilution) and 4′,6-diamidino-2-phenylindole (DAPI) staining. Plates were imaged on a Cytation7 (Agilent), and foci were counted using Gen5 software. The 50% endpoint neutralization titers were determined as the reciprocal of the highest dilution providing ≤ half of the number of foci obtained from the negative control well (plain DMEM mixed with 100 PFU virus).

### Histopathology Analyses

Procedures as described previously (24, 25). Both mouse and hamster lung tissues were fixed in 10% neural buffered formalin overnight and then processed for paraffin embedding. The 5-μm sections were stained with hematoxylin and eosin for histopathological examinations. Images were scanned using an Aperio ImageScope. Blinded samples were graded by a licensed pathologist for the following 14 categories: % lung section affected, alveolar wall thickening, alveolar airway infiltrates, perivascular infiltrates, perivascular edema, alveolar osteoid foci, type II pneumocyte hyperplasia, atypical pneumocyte hyperplasia, alveolar metaplasia, bronchiolar necrosis, mucosal hyperplasia, bronchiole airway infiltrates, proteinaceous fluid, vasculitis.

Lesion scoring: 0 = none, 1 = minimal, 2 = mild, 3 = moderate, 4 = severe. A graph was prepared by summing up the score in each category. The following antibodies were used for immunohistochemistry (IHC) staining: rabbit anti-CD8 alpha (cytotoxic T cell) (#98941S, Cell Signaling Technology, USA) (1:250 dilution) and rabbit anti-CD4 (helper T cell) (#25229S, Cell Signaling Technology, USA) (1:50 dilution).

### Statistical analysis

One-way or two-way ANOVA, or paired t test was used to calculate statistical significance through GraphPad Prism (10.0.0) software for Windows, GraphPad Software, San Diego, California USA, www.graphpad.com.

## Author Contributions

P. S., C. B. S., and T. T. W. conducted the experiments, analyzed the data, and wrote the paper. S.L. made live attenuated vaccine candidate viruses and K.S assisted in animal experiments & qRT-PCR.

